# The *Klebsiella pneumoniae ter* operon enhances stress tolerance

**DOI:** 10.1101/2022.12.02.518861

**Authors:** Sophia Mason, Jay Vornhagen, Sara N. Smith, Laura A. Mike, Harry L.T. Mobley, Michael A. Bachman

**Affiliations:** Department of Pathology, Michigan Medicine, University of Michigan, Ann Arbor, United States of America; Department of Microbiology & Immunology, Michigan Medicine, University of Michigan, Ann Arbor, United States of America; Department of Medical Microbiology & Immunology, University of Toledo, Toledo, United States of America

## Abstract

Healthcare-acquired infections are a leading cause of disease in patients that are hospitalized or in long-term care facilities. *Klebsiella pneumoniae* (Kp) is a leading cause of bacteremia, pneumonia, and urinary tract infections in these settings. Previous studies have established that the *ter* operon, a genetic locus that confers tellurite oxide (K_2_TeO_3_) resistance, is associated with infection in colonized patients. Rather than enhancing fitness during infection, the *ter* operon increases Kp fitness during gut colonization; however, the biologically relevant function of this operon is unknown. First, using a murine model of urinary tract infection, we demonstrate a novel role for the *ter* operon protein TerC as a bladder fitness factor. To further characterize TerC, we explored a variety of functions, including resistance to metal-induced stress, resistance to ROS-induced stress, and growth on specific sugars, all of which were independent of TerC. Then, using well-defined experimental guidelines, we determined that TerC is necessary for tolerance to ofloxacin, polymyxin B, and cetylpyridinium chloride. We used an ordered transposon library constructed in a Kp strain lacking the *ter* operon to identify genes required to resist K_2_TeO_3_− and polymyxin B-induced stress, which suggested that K_2_TeO_3_-induced stress is experienced at the bacterial cell envelope. Finally, we confirmed that K_2_TeO_3_ disrupts the Kp cell envelope, though these effects are independent of *ter*. Collectively, the results from these studies indicate a novel role for the *ter* operon as stress tolerance factor, therefore explaining its role in enhancing fitness in the gut and bladder.

## Introduction

*Klebsiella pneumoniae* (Kp) is a pathogenic member of the *Enterobacteriaceae* family (Order *Enterobacterales*) and a cause of pneumonia, UTI, and bloodstream infections (1). Importantly, Kp has a high potential for antibiotic resistance and is the third leading cause of global deaths attributable to pathogens with the potential for antibiotic resistance (2). Infection with antibiotic-resistant Kp is associated with high mortality rates, with mortality rates ranging from approximately 20-40% (3–5). Multiple studies have demonstrated a strong association (odds ratio ∼4.0) between gut colonization and infection, indicating that infectious Kp originates from the gut of colonized patients (6–9). Colonization rates are variable, ranging as high as >75% in hospitalized patients (10), though several large studies indicate a colonization rate closer to 20% (6–8), depending on seasonal effects (11). Therefore, Kp can successfully and silently tolerate the hostile gut environment before causing disease (12). More information about the factors that influence Kp colonization and infection is necessary given the high risk of infection posed to patients colonized by Kp, especially in the context of increasing levels of antibiotic resistance.

Our recent work revealed that the presence of the Kp *ter* operon is associated with bacteremia and pneumonia in colonized patients (13). Experimental interrogation of the *ter* operon using an isogenic mutant revealed that the *ter* operon increases Kp fitness during gut colonization rather than conferring a fitness advantage during bacteremia and pneumonia (13, 14). In particular, a *terC* mutant is less fit in mice with an increased abundance of bacteria known to produce short-chain fatty acids (SCFA). SCFAs, specifically acetate, can inhibit the growth of Kp in a pH-dependent manner (15). However, exogenous administration of SCFAs to mice during gut colonization, but not *in vitro*, results in a fitness defect dependent on the *ter* operon. Therefore, the inhibitory effect of SCFAs is dependent on the presence of specific indigenous gut microbiota (14). This suggests an alternative biological role for the *ter* operon during Kp pathogenesis, wherein this operon is responsive to a yet-undefined stresses.

The molecular function of the *ter* operon is cryptic. This operon confers resistance to the oxyanion form of the rare non-essential trace element tellurium, tellurite oxide (TeO_3_^−2^). Additionally, the mechanism by which TeO_3_^−2^ damages bacterial cells is unclear. Tellurium is occasionally grouped with other transition metals, which exhibit antibacterial properties; however, as a chalcogen, its toxicity likely differs from that of transition metals. Furthermore, TeO_3_^−2^ is largely absent in the human body or medical settings (16, 17). Therefore, physiologically relevant stress or stresses other than TeO_3_^−2^ must explain the strong association between the *ter* operon and Kp pathogenesis. The biochemical mechanisms of TeO_3_^−2^-induced stress affect several critical pathways and therefore induces pleiotropic stress (reviewed in (18)), making it challenging to infer what physiologically relevant stresses interact with the *ter* operon. Thought to be primarily responsible for its toxicity, the strong oxidizing ability of TeO_3_^−2^ has several secondary adverse consequences for the bacterial cell (19). The reduction of TeO_3_^−2^ creates hydroxyl radicals by inhibiting heme biosynthesis (20). As a result, these hydroxyl radicals abrogate DNA synthesis and protein synthesis, exhaust cellular reductases, and oxidize membrane lipids (21–24). In conjunction with the proposed biochemical mechanism of TeO_3_^−2^-induced stress, the *ter* operon may play a role in resistance to colicins (family A, B, and K) and bacteriophage (16). Notably, many of the biochemical mechanisms of TeO_3_^−2^-induced stress are akin to those that Kp may encounter during its pathogenesis. For example, in the gut where the *ter* operon is conditionally required for complete fitness, colicins are important weapons of bacterial warfare, and host antimicrobial peptides limit pathogen proliferation (25, 26). These stresses kill by attacking the bacterial membrane and DNA similar to the effects of TeO_3_^−2^ (25, 26). Therefore, exploring the molecular function of the *ter* operon is likely to reveal interesting facets of Kp pathogenesis.

Here, we interrogate the physiologically relevant role of the Kp *ter* operon and associated TeO_3_^−2^-induced stress. First, we identified a novel role for the *ter* operon as a fitness factor during urinary tract infection (UTI). Concordant with a fitness impact in multiple body sites, we found that the *ter* operon is involved in stress tolerance, a phenotype wherein more tolerogenic cells die slower than less tolerogenic cells in the presence of harmful agents. This phenotype comports with a general, rather than a specific, stress response. Finally, using a systematic approach, we identified novel genes associated with TeO_3_^−2^ resistance in Kp lacking the *ter* operon and indeed found that TeO_3_^−2^ acts on diverse biological pathways, which corresponds to a need for a general mechanism of stress response. Collectively, these data suggest a role for the *ter* operon in responding to envelope destabilization, resulting in enhanced stress tolerance and potentially explaining how *ter* operon function enhances fitness during UTI and gut colonization.

## Results

### TerC is a bladder fitness factor during urinary tract infection

Our previous study in which we identified the strong association between the *ter* operon and infection was limited to patients that developed pneumonia and bacteremia (13). However, Kp is also an important cause of UTI in colonized patients (7, 11). In a previous survey of 2,549 urine associated Kp isolates, we found that 8.4% contained *ter* (14). To test the role of the *ter* operon in UTI, we competed our *terC* mutant and wildtype (NTUH-K2044) strains 1:1 in a well-established murine transurethral infection model (27). Deletion of *terC* is sufficient to confer susceptibility to K_2_TeO_3_ and is complemented by *terZ-F* (13, 14). Here, we observed a 2-log fitness defect for the *terC* mutant strain in the bladder of infected mice (Fig. 1). Furthermore, the *terC* mutant strain was mostly absent from the bladder of infected mice (only 2/17 with detectable CFU). The observed bladder fitness defect was not due to differences in growth in urine (Fig. S1A-C) and was not observed *ex vivo* in bladder homogenate (Fig. S1D), indicating that the whole organism, or at least viable tissue is required to observe a TerC-dependent fitness defect. The finding that the *ter* operon was required for complete fitness in the bladder *in vivo* was somewhat surprising, as we have previously demonstrated that TerC is required for complete fitness in the gut in a microbiome-dependent manner (14) and bladder-indigenous microbiota are either sparse or absent (28). Therefore, these findings suggest a function for the *ter* operon that explains a role for fitness in both the gut and bladder.

**Figure 1.**
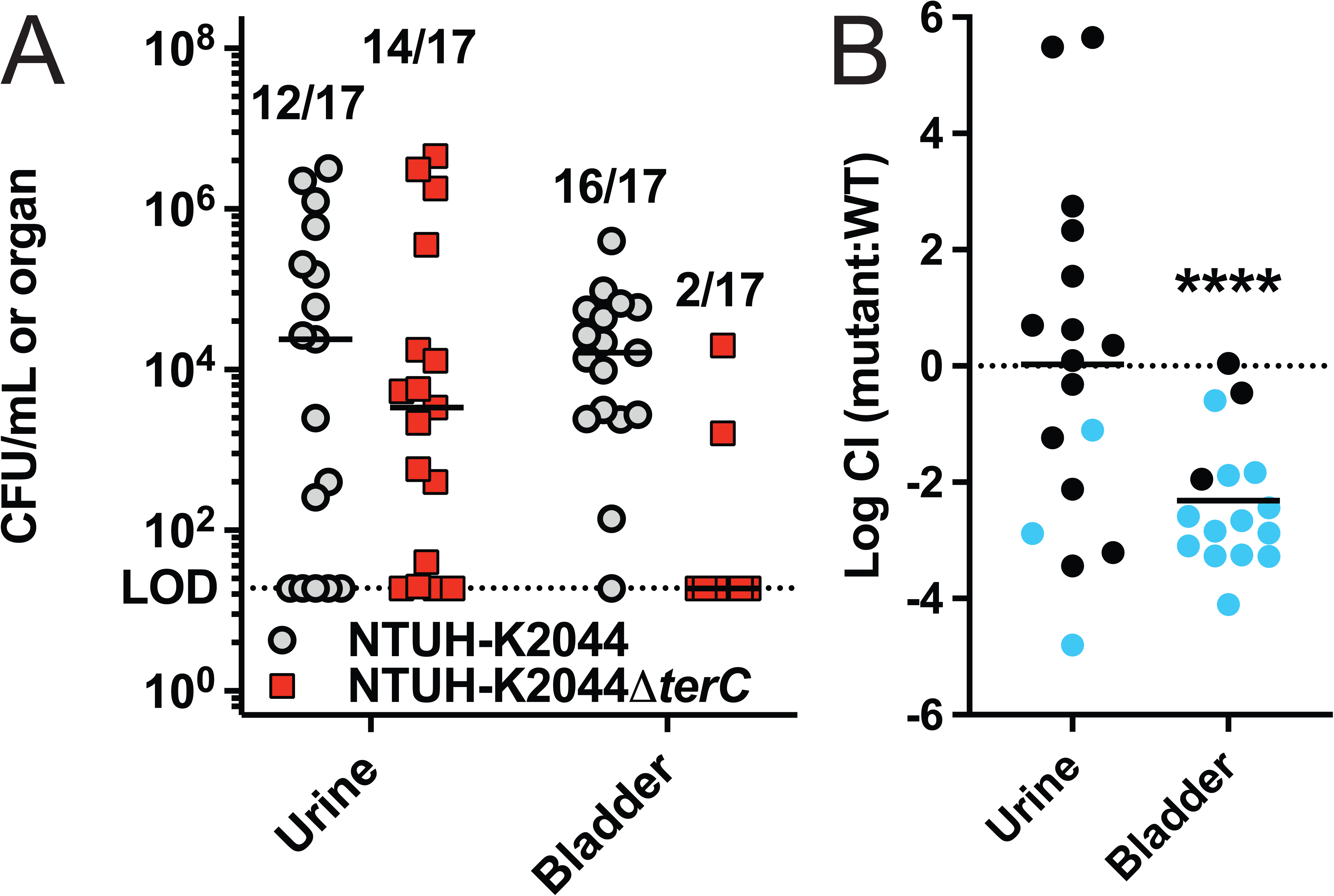
TerC is necessary for complete fitness in the bladder during urinary tract infection. Mice were transurethrally inoculated with approximately 10^8^ CFU of a 1:1 mix of WT NTUH-K2044 and NTUH-K2044Δ*terC*. (A) Bacterial burden in the urine and bladder was measured after 48 hours, and (B) log_10_ competitive index (CI) of the mutant strain compared to the WT strain was calculated for each organ (N = 17, mean displayed, ****P < 0.00005, one-sample *t*-test). From 20 inoculated mice, CFU were recovered from 17 mice. The numbers above each column in A indicate the number of mice (of 17) with detectable CFU. Blue circles had no recoverable NTUH-K2044Δ*terC*. CI was calculated using the limit of detection CFU value when no CFU were recovered.

### TerC is not required for growth on common sugars

Given the finding that TerC is necessary for complete fitness in two dissimilar body sites, we turned to a hypothesis-driven approach to identify the mechanisms that unify these phenotypes. Structure-based prediction of the function suggested that TerC may act as a sugar transporter (14). Specifically, the highest-confidence functional predictions were hexose:proton symporter activity, galactose transmembrane transporter activity, arabinose transmembrane transporter activity, and fucose transmembrane transporter activity. This activity could explain the fitness defect observed in multiple, unrelated body sites, as perturbations in metabolic flexibility are known to impact Kp site-specific fitness, including the gut (29). To test this hypothesis, we compared growth of WT and *terC* mutant strains containing an empty pACYC184 vector (NTUH-K2044 pVector and NTUH-K2044Δ*terC* pVector, respectively) and a *terC* complement strain expressing *terZ-F in trans* (NTUH-K2044Δ*terC* pTerZ-F) in M9 minimal medium supplemented with various sugars at a final concentration of 0.5%. We previously demonstrated that expression of *terZ-F* is required to complement the K_2_TeO_3_ resistance phenotype (14). TerC was dispensable for growth in the presence of all sugars tested (Figure S2), which include: the hexoses fucose, galactose, glucose, and rhamnose; the pentoses arabinose and xylose; and the disaccharides lactose and sucrose. The NTUH-K2044Δ*terC* pTerZ-F strain had small but significant growth defects in lactose, sucrose, and rhamnose, and grew significantly better than NTUH-K2044 pVector in galactose based on area under the curve analysis (Figure S2). Although not exhaustive, these data indicate that TerC is not required for growth on these eight common sugars.

### TerC is dispensable for resistance to metal and radical oxygen species-induced stress

We next hypothesized that the *ter* operon may be involved in resistance to transition metal-induced stress. Metal-responsive genes are important fitness factors during UTI (30, 31) and gut colonization (32). Moreover, several transition metal resistance operons are co-localized on *ter* operon containing plasmids (14), suggesting a conserved role for transition metal resistance for these plasmids. To test this hypothesis, we determined the minimum inhibitory concentration (MIC) of several first-row transition metals with known function as micronutrients and that are toxic in excess through induction of redox stress (reviewed in (33) and (34)). As expected, the *terC* mutant strain was more susceptible to K_2_TeO_3_ than the WT strain, and this phenotype was complemented by expressing *terZ-F in trans* (Figure S3A). No differences in MIC were observed for any transition metals, indicating that the *ter* operon is dispensable for resistance to transition metal-induced stress (Figure S3A). Moreover, the MIC of K_2_TeO_3_ was substantially lower than that of the tested first-row transition metals, indicating that the biological activity of the first-row transition metals differs from that of K_2_TeO_3_.

As K_2_TeO_3_ is a potent generator of radical oxygen species (ROS), next we explored the hypothesis that the *ter* operon may be involved in resistance to ROS-induced stress (18, 20). Like metal-induced stress, resistance to ROS has been implicated in bacterial fitness during UTI (35, 36) and gut colonization (37). Moreover, some studies have suggested that the *ter* operon is under transcriptional control by OxyR (38, 39). To test this hypothesis, we determined the impact of TerC on the MIC of three cell-permeable ROS generators: hydrogen peroxide (H_2_O_2_), and the superoxide-generating redox-cycling compounds paraquat and menadione. As above, TerC was necessary for K_2_TeO_3_ resistance; however, TerC was dispensable for resistance to H_2_O_2_, paraquat, and menadione (Figure S3B). Collectively, these data suggest an alternative function for the *ter* operon.

### TerC plays a role in stress tolerance

It is perhaps not surprising that the *ter* operon is dispensable for metal and ROS-induced stress, as there are several factors, such as superoxide dismutase, glutathione, and catalase (reviewed in (40)), that play an important role in detoxifying ROS. Correspondingly, metal and ROS resistance genes are important fitness factors during Kp lung infection (41, 42) where the *ter* operon is dispensable (13). It may be the case that the *ter* operon plays a different role in resisting TeO_3_^−2^ and the pleiotropic stress it induces. Therefore, we further considered the structure-function relationship of TerC. While TerC does not appear to be a dedicated sugar transporter, it has been suggested that Ter proteins form a stress-sensing membrane complex anchored by TerC (43). A potential role for TerC in sensing or responding to envelope stress was intriguing, as maintenance of the cell envelope is critical for stress tolerance (44, 45). Notably, TerC is membrane-bound (46).

To explore a role for TerC and the *ter* operon in stress tolerance, we turned to the definitions and guidelines for research on antibiotic persistence (47). These guidelines outline the experimental approach for differentiating stress resistance, tolerance, and persistence. Briefly, resistance is defined as difference in the MIC of a stress, whereas tolerance and persistence are defined as enhanced survival during stress exposure with no change in MIC. Rather, tolerant cells die more slowly during stress exposure, and persister cells are a small subpopulation of tolerant cells that survive stress exposure better than the general population. Mechanistically, tolerance is often mediated by specific, population-wide stress responses (reviewed in (48)), whereas persister cells are a stochastically formed sub-population (reviewed in (49)). These phenotypes can be differentiated by measuring the kill curve in the presence of stress and summarized as the minimum duration of killing at a pre-defined percentage of the initial population (MDK_%_). Chemical concentrations well above (≥10X MIC) are used for these experiments. Differences in stress tolerance between two populations are defined by differential killing of a large fraction of the initial population, such as differences in 90% (MDK_90_) or 99% (MDK_99_). The difference in persistence between two populations is defined by a biphasic kill curve, wherein initial killing occurs as a similar rate (*e.g.*, identical MDK_90_ or MDK_99_), then transitions to a differential biphasic state, such as differences in 99.9% (MDK_99.9_) or 99.99% (MDK_99.99_) of the initial population.

Using these guidelines, we assessed the role of TerC in stress tolerance. First, we characterized the dynamics of K_2_TeO_3_-mediated killing and observed more rapid killing of the *terC* mutant compared to its parent strain (Figure 2Ai-ii), which was determined to be significantly different based on AUC analysis (Figure 2Aiii) and interpolation of exact MDK values (Figure 2Aiv). Finally, TerC-dependent survival at 4 hours post stress exposure was complemented *in trans* (Figure 2Av). Next, we explored killing dynamics in response to ofloxacin-induced stress. Fluoroquinolones, such as ofloxacin, kill bacterial cells through inhibition of DNA gyrase and are commonly used to generate persister cells in a laboratory setting (50). The MIC of ofloxacin was identical between *terC* mutant and its parent strain (Figure 2Bi), indicating TerC is not an ofloxacin resistance factor. Interestingly, we observed significant differences in the kill curves between these two strains (Figure 2Bii-iii). Interpolation of MDK values revealed a significant difference between the MDK_90_ and MDK_99_ of the *terC* mutant and its parent (Figure 2Biv), indicating that the wildtype strain was significantly more tolerant to ofloxacin-induced stress than the *terC* mutant. The MDK_99.9_ and MDK_99.99_ were incalculable for the wildtype strain, as ofloxacin failed to kill this proportion of cells. As above, TerC-dependent survival at 4 hours post stress exposure was complemented *in trans* (Figure 2Bv). Collectively, these data suggest that TerC may be necessary for ofloxacin tolerance.

**Figure 2.**
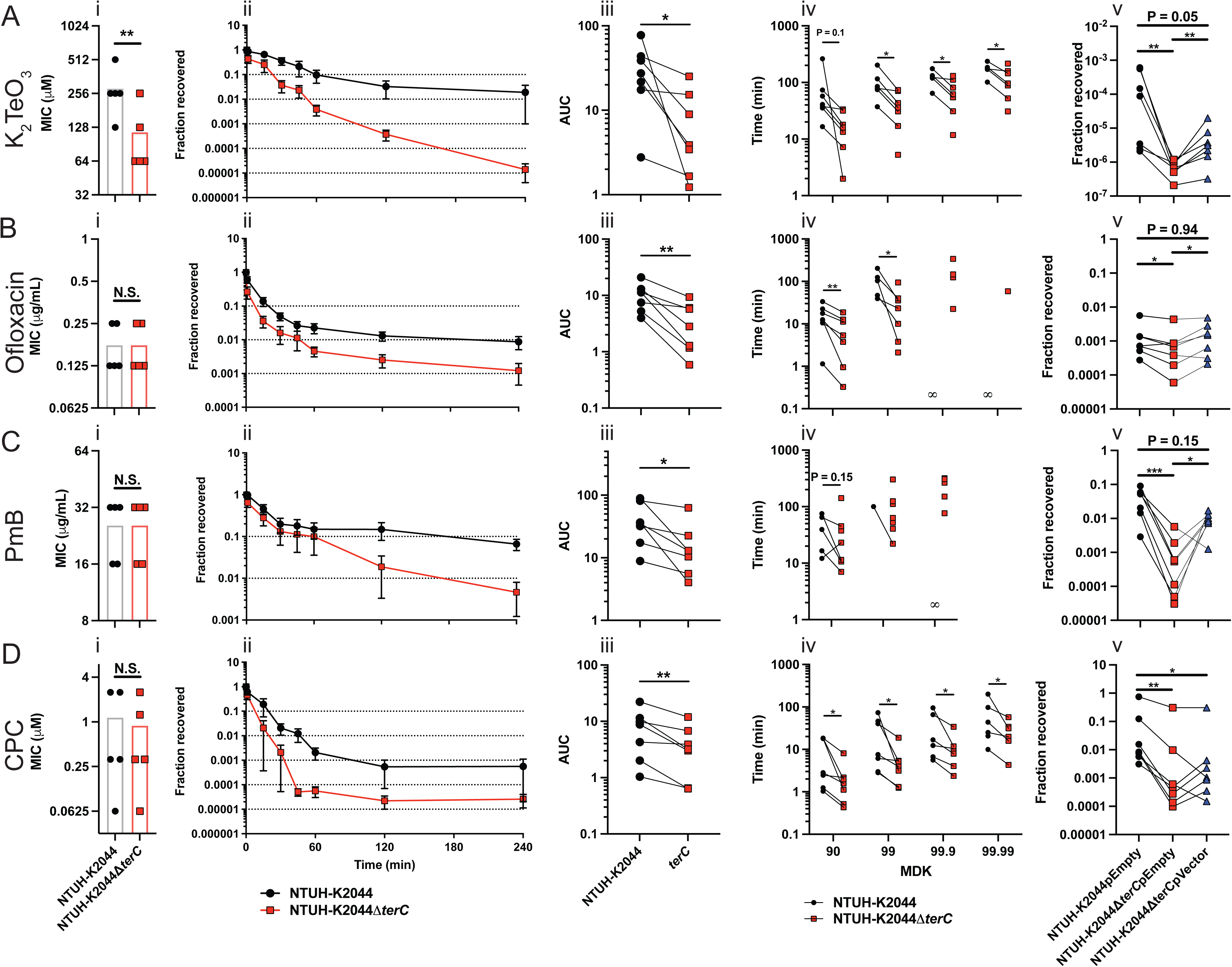
TerC is necessary for tolerance to several stresses. The (A) K_2_TeO_3_ minimum inhibitory concentration (i, n = 5 independent experiments, median displayed, **P < 0.005, ratio paired *t*-test) was calculated for NTUH-K2044 and NTUH-K2044Δ*terC* strains using broth microdilution. Kill curves (ii, n = 7 independent experiments, mean displayed ± SEM) were generated for these strains using by adding a standard K_2_TeO_3_ concentration of 1 mM to overnight cultures and (iii) AUC and (iv) MDK were calculated from these kill curves (n = 7 per group, *P < 0.05, **P < 0.005, ratio paired *t*-test, ∞ indicates an incalculable MDK). Finally, NTUH-K2044 pVector, NTUH-K2044Δ*terC* pVector, and NTUH-K2044Δ*terC* pTerZ-F survival was assessed at 4 hours (240 minutes) post 1 mM K2TeO3 exposure (v, n = 6-7 independent experiments, median displayed, *P < 0.05, **P < 0.005, ***P < 0.0005, one-way ANOVA followed by Tukey’s multiple comparison test). These experiments were repeated for ofloxacin (B), polymyxin B (PmB, C), and cetylpyridinium chloride (CPC, D). The standard concentrations for killing assays for ofloxacin, polymyxin B, and cetylpyridinium chloride were 250 μg/mL, 500 μg/mL, and 25 μM, respectively. For iii, iv, and v, connecting lines indicate paired biological replicates.

To investigate this tolerance phenotype further, we tested a second antibiotic, polymyxin B (PmB), which has a different mechanism of action than ofloxacin: disruption of the bacterial cell envelope. As was observed with ofloxacin, the MIC of PmB was identical for the *terC* and its parent strain (Figure 2Ci). Notably, we observed significant differences in the kill curves between these two strains in response to PmB-induced stress, wherein the wildtype strain was killed more slowly that the *terC* mutant (Figure 2Cii-iii). The MDK_90_ of the *terC* mutant trended lower than its parent strain and the MDK_99_ and MDK_99.9_ was incalculable for the wildtype strain (Figure 2Civ). Complementation *in trans* restored survival at 4 hours post stress exposure (Figure 2Cv). As with ofloxacin, these data indicate that the *terC* mutant is less tolerant to PmB-induced stress than its parent strain. Finally, we repeated these experiments with cetylpyridinium chloride (CPC), a quaternary ammonium compound that disrupts the cell envelope, leading to leakage of intracellular contents and death (51). As with PmB, no differences between the MIC of the wildtype strain at the *terC* mutant was observed (Figure 2Di); however, the *terC* mutant was killed more rapidly that its parent strain due to CPC-induced stress, which was reflected in all MDK values (Figure 2Dii-iv). Finally, 4-hour post stress exposure survival was complemented *in trans* (Figure 2Dv). Collectively, these data indicate that TerC is necessary for tolerance to stresses with distinct mechanisms of action and potentially explains the requirement of TerC for complete fitness in the gut and bladder.

### A systematic screen reveals diverse genes associated with K_2_TeO_3_ resistance

The finding that TerC is required for tolerance, but not resistance, to stresses with differing mechanisms of action comports with its requirement for resistance to the pleiotropic stress induced by K_2_TeO_3_; however, this finding does not implicate a specific pathway or cellular compartment of action for the *ter* operon. To determine if K_2_TeO_3_ acts on a specific pathway or cellular compartment, we turned to a systematic approach to identify genes and pathways required to mitigate the effects of the K_2_TeO_3_-induced stress when the *ter* operon was absent. To this end, we employed an ordered, condensed transposon (Tn) library constructed in the Kp strain KPPR1, which doesn’t encode the *ter* operon and is susceptible to K_2_TeO_3_ (Figure 3A-B). This Tn library contains individual insertions in 3,733 genes, covering 72% of all available open reading frames in the genome (52). First, we determined the concentration at which KPPR1 growth was partially inhibited by K_2_TeO_3_ to be approximately 1 μM. We aimed to screen our Tn library at this concentration to identify mutants conferring both resistance (increase in growth) and susceptibility (decrease in growth) to K_2_TeO_3_. We then cultured each mutant in the presence of 1 μM K_2_TeO_3_ and assessed growth by measuring growth at OD_600_ (Figure 3C). If the growth of a mutant was two standard deviations above or below the mean growth of all mutants, we categorized the interrupted gene as potentially being important for K_2_TeO_3_ resistance. Initially, we found 129 candidate susceptible mutants and 15 candidate resistant mutants (Figure 3C). It was not surprising that we identified more susceptible than resistant mutants, as previous studies using other bacteria to identify genes involved in K_2_TeO_3_ resistance required several passages in the presence of K_2_TeO_3_ to identify such loci (20). As our screen was performed at a standard concentration of K_2_TeO_3_, we next aimed to validate our candidate mutants. We preliminarily excluded 14 mutants due to an observed growth defect in LB in the absence of K_2_TeO_3_. We then determined the concentration of K_2_TeO_3_ that inhibited 50% of bacterial growth (IC_50_) for our remaining candidates. Of the 138 candidates, 79 had a K_2_TeO_3_ IC_50_ that corroborated our original screen, and 29 had a significantly lower (q ≤ 0.1) IC_50_ value than that of the parent strain (Table 1, Figure 3D, Table S1). To further characterize these genes associated with K_2_TeO_3_ resistance, we assessed the gene function using the Kyoto Encyclopedia of Genes and Genomes (KEGG) (Figure 3E, Table S1). Metabolism (n = 12/29) and “Other” (n = 12/29) were the most well-represented categories. Assignment of more granular function of these genes revealed that the most common function was carbohydrate metabolism (n = 5/29), followed by genetic information processing (n = 4/29). These findings suggest that many diverse genes are necessary for growth in the presence of K_2_TeO_3_, rather than a set of related genes with conserved function.

**Figure 3.**
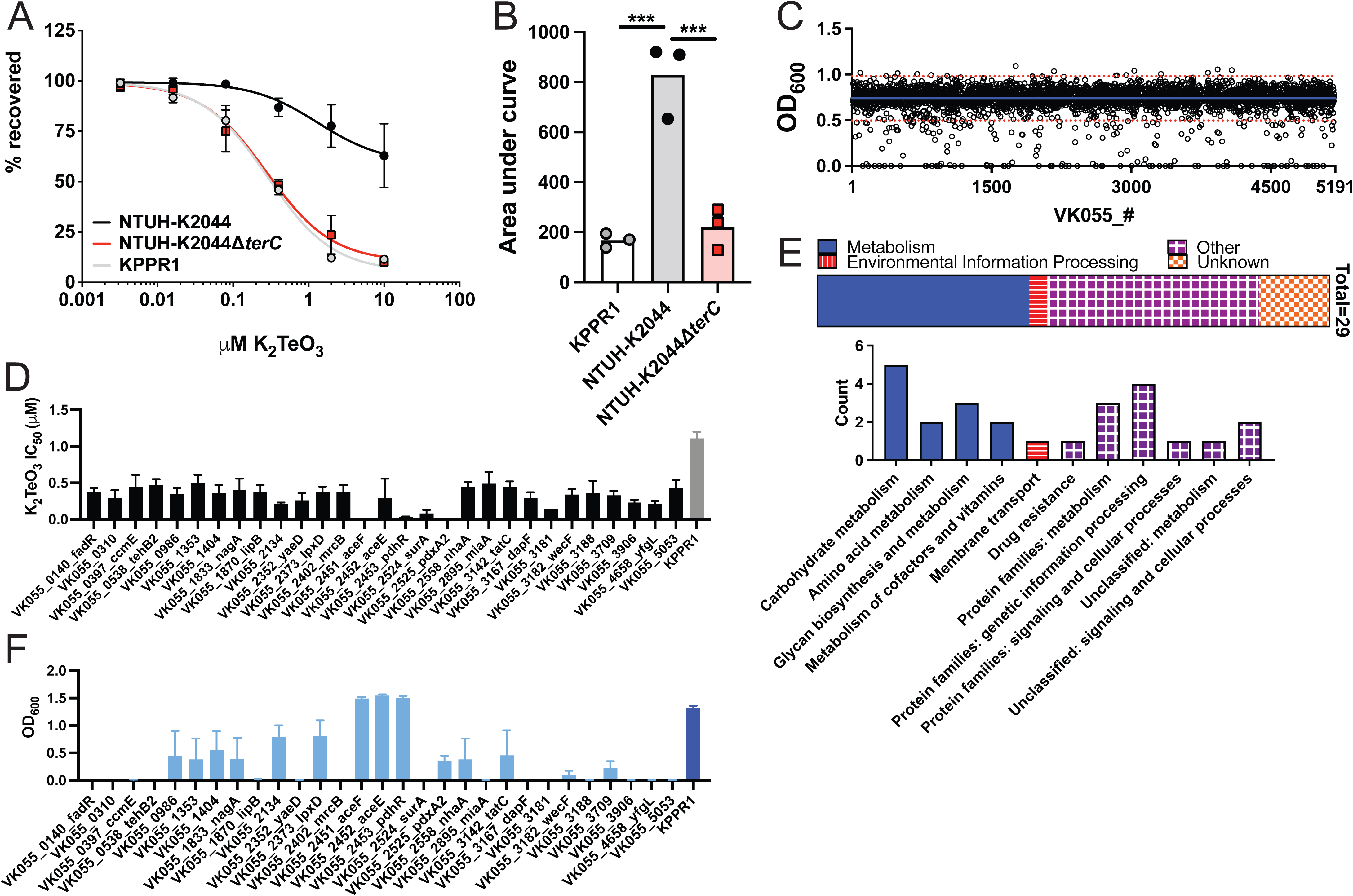
Systematic screen of K_2_TeO_3_ resistance. The KPPR1 strain, which lacks the ter operon, and the NTUH-K2044 and NTUH-K2044Δ*terC* strains were cultured in increasing concentrations of K_2_TeO_3_ (A) and area under the curve (AUC) was calculated from these dose-response curves (B, mean displayed ± SEM, ****P < 0.0005, one-way ANOVA followed by Tukey’s multiple comparison test). (C) 3,733 individual Tn insertion mutants were cultured in the presence of 1 μM K_2_TeO_3_ and measured at OD_600_. The blue line is the mean OD_600_ and the red lines are ± 2 S.D. from the mean. Each symbol is an individual mutant, ordered by their gene number (VK055_#). (D) Exact K_2_TeO_3_ IC_50_ values of validated Tn insertion mutants (n = 3-5 independent experiments). (E) KEGG BRITE Functional Hierarchies were assigned for validated Tn insertion mutants. The highest order hierarchies are shown in the horizontal stacked bar chart, and the second highest order hierarchies are shown in the unstacked bar chart. (F) Validated Tn insertion mutants were cultured in the presence 0.5 µg/mL PmB (n = 3 independent experiments).

**Table 1.** Insertion mutants involved in K_2_Te0_3_ resistance.

| Gene Name | Annotation | Subcellular Location | K<sub>2</sub>TeO<sub>3</sub> IC<sub>50</sub>±SEM (KPPR1 = 1.11±0.09) | P Value | q Value |
| --- | --- | --- | --- | --- | --- |
| VK055_0140_fadR | fatty acid metabolism transcriptional regulator FadR | cytoplasm | 0.37±0.06 | 0.003060 | 0.010447 |
| VK055_0310 | bacterial regulatory helix-turn-helix, lysR family protein |  | 0.29±0.11 | 0.000093 | 0.000656 |
| VK055_0397_ccmE | cytochrome c-type biogenesis protein CcmE | integral component of membrane | 0.44±0.17 | 0.002301 | 0.009113 |
| VK055_0538_tehB2 | tellurite resistance protein TehB | cytoplasm | 0.47±0.08 | 0.006082 | 0.019425 |
| VK055_0986 | hypothetical protein | N/A | 0.35±0.08 | 0.001477 | 0.006521 |
| VK055_1353 | putative L,D-transpeptidase YcfS | periplasmic space | 0.50±0.11 | 0.027590 | 0.068284 |
| VK055_1404 | periplasmic glucans biosynthesis protein MdoG | periplasmic space | 0.36±0.11 | 0.001024 | 0.005068 |
| VK055_1833_nagA | N-acetylglucosamine-6-phosphate deacetylase NagA | cytoplasm | 0.40±0.16 | 0.002208 | 0.009106 |
| VK055_1870_lipB | lipoyl(octanoyl) transferase LipB | cytoplasm | 0.38±0.09 | 0.002789 | 0.009860 |
| VK055_2134 | putative dTDP-glucose pyrophosphorylase |  | 0.21±0.02 | 0.000011 | 0.000120 |
| VK055_2352_yaeD | D,D-heptose 1,7-bisphosphate phosphatase GmhB | cytoplasm | 0.26±0.10 | 0.000020 | 0.000176 |
| VK055_2373_lpxD | UDP-3-O-[3-hydroxymyristoyl] glucosamine N-acyltransferase LpxD | cytoplasm | 0.37±0.08 | 0.002434 | 0.009268 |
| VK055_2402_mrcB | penicillin-binding protein 1B MrcB | peptidoglycan-based cell wall | 0.38±0.09 | 0.002789 | 0.009860 |
| VK055_2451_aceF | pyruvate dehydrogenase E2 component AceF | cytoplasm | 0.01±0.00 | 0.000001 | 0.000001 |
| VK055_2452_aceE | pyruvate dehydrogenase E1 component AceE |  | 0.29±0.27 | 0.000001 | 0.000001 |
| VK055_2453_pdhR | transcriptional repressor for pyruvate dehydrogenase complex PdhR | cytoplasm | 0.03±0.01 | 0.000001 | 0.000001 |
| VK055_2524_surA | peptidyl-prolyl cis-trans isomerase SurA | periplasmic space | 0.08±0.05 | 0.000001 | 0.000001 |
| VK055_2525_pdxA2 | 4-hydroxythreonine-4-phosphate dehydrogenase PdxA2 | cytoplasm | 0.02±0.00 | 0.000001 | 0.000001 |
| VK055_2558_nhaA | na<sup>+</sup>/H<sup>+</sup> antiporter NhaA | integral component of membrane | 0.45±0.06 | 0.015340 | 0.039965 |
| VK055_2895_miaA | tRNA dimethylallyltransferase MiaA | cytoplasm | 0.49±0.16 | 0.013757 | 0.038914 |
| VK055_3142_tatC | sec-independent protein translocase protein TatC | integral component of membrane | 0.45±0.07 | 0.014802 | 0.039606 |
| VK055_3167_dapF | diaminopimelate epimerase DapF | cytoplasm | 0.29±0.08 | 0.000148 | 0.000977 |
| *VK055_3181* | enterobacterial common antigen polymerase WzyE | integral component of membrane | 0.14±0.00 | 0.000001 | 0.000002 |
| *VK055_3182_wecF* | dTDP-N-acetylfucosamine:lipid II N-acetylfucosaminyltransferase WecF | integral component of membrane | 0.34±0.07 | 0.001236 | 0.005826 |
| *VK055_3188* | UDP-N-acetyl-D-mannosaminuronic acid dehydrogenase WecC |  | 0.36±0.17 | 0.000323 | 0.001880 |
| *VK055_3709* | shikimate kinase AroK | cytoplasm | 0.33±0.06 | 0.000970 | 0.005055 |
| *VK055_3906* | hypothetical protein | N/A | 0.23±0.04 | 0.000021 | 0.000177 |
| *VK055_4658_yfgL* | outer membrane protein assembly factor BamB | integral component of membrane | 0.21±0.04 | 0.000007 | 0.000088 |
| *VK055_5053* | DNA gyrase inhibitor SbmC | cytoplasm | 0.43±0.11 | 0.008174 | 0.025287 |

To determine if genes involved in K_2_TeO_3_ resistance are also involved in resistance to stressors where TerC was required for tolerance, we screened mutants validated for increased K_2_TeO_3_ sensitivity for growth in the presence of PmB. Interestingly most genes involved in K_2_TeO_3_ resistance were also required for growth in the presence of PmB (Figure 3F). The pyruvate dehydrogenase complex (*aceF*, *aceE*, *pdhR*) was one notable exception to this finding. Given that PmB is a membrane-active antibiotic, this finding further supports the indication that K_2_TeO_3_ destabilizes the Kp envelope. Moreover, this suggests that the function of the *ter* operon is to aid in envelope stabilization or respond to envelope destabilization, leading to enhanced stress tolerance, and enhanced fitness in the gut and bladder.

Gene ontology biological process enrichment analysis revealed a single enriched pathway amongst these individual genes: enterobacterial common antigen (ECA) biosynthetic process (42.23-fold enrichment, FDR P value = 5.25 x 10^−2^). Notably, additional ECA genes *wecG*, *wecA* narrowly missed our validation criteria (Table S1). This finding supports a potential role for the *ter* operon in maintaining envelope stability or responding to envelope destabilization, which comports with a role in stress tolerance.

### K_2_TeO_3_ disrupts the Kp cell envelope

Given that the finding that many genes associated with K_2_TeO_3_ resistance in a strain lacking *ter* may play a role in maintaining envelope stability we next aimed to determine if K_2_TeO_3_ disrupts the cell envelope, and if the *ter* operon stabilizes the envelope. Fluorescence-based Ethidium bromide (EtBr) accumulation assays can be used to assess envelope damage (53, 54), wherein EtBr accumulates in the periplasm or intercalates in cellular DNA following cell envelope disruption. As colistin has been shown to disrupt both the outer and inner membrane of the Gram-negative envelope (55), we used PmB as a positive control. Following exposure to K_2_TeO_3_ and PmB at the same concentrations used in killing assays (Figure 2), Kp displayed higher levels of EtBr accumulation than that of the no treatment controls (Figure 4). This phenotype was independent of TerC, suggesting that the effects of *ter* are downstream of initial disruption of the cell envelope (Figure 4). These data demonstrate that K_2_TeO_3_ disrupts the cell envelope and in conjunction with the data in Figure 2Aii (difference in fraction recovered at 60 minutes post-K_2_TeO_3_ exposure) suggest that the effects of *ter* are downstream of envelope destabilization.

**Figure 4.**
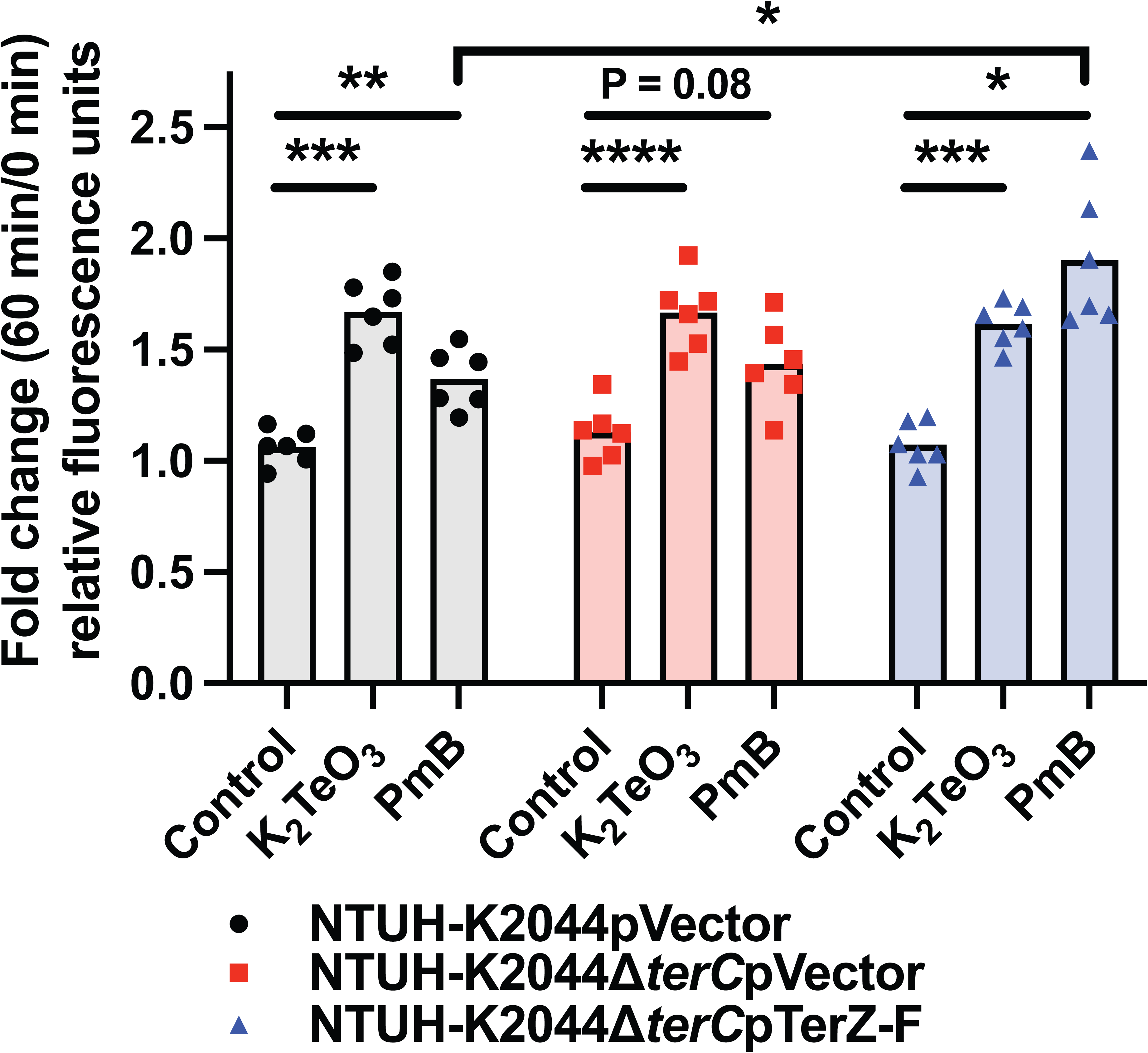
EtBr accumulates in K_2_TeO_3_ and polymyxin B treated Kp. Stationary phase NTUH-K2044 pVector, NTUH-K2044ΔterC pVector, and NTUH-K2044ΔterC pTerZ-F were exposed to 1 mM K_2_TeO_3_ or 500 μg/mL PmB. EtBr accumulation (fluorescence) was measured at baseline and at one-hour post-exposure (n = 6 independent experiments, mean displayed, *P < 0.05, **P < 0.005, ***P < 0.0005, one-way ANOVA followed by Tukey’s multiple comparison test).

## DISCUSSION

The work presented here advances our understanding of the physiological role of the Kp *ter* operon. Our results indicate that a physiological role of the *ter* operon is to respond to envelope stress during colonization and infection of specific body sites to tolerate these stressful environments and enhance fitness. These effects are likely downstream of the initial insult to the envelope. Previously, we have shown that this cryptic operon is highly associated with Kp pneumonia and bacteremia in colonized patients (13), and further work demonstrated that this association was due to a TerC-dependent fitness advantage conferred in the gut (14). Here, we demonstrate a novel role for TerC as a fitness factor during urinary tract infection. The identification of TerC as a bladder fitness factor using a hypervirulent Kp strain (NTUH-K2044) is noteworthy, as several recent reports have indicated that hvKp are an important cause of asymptomatic bacteriuria and UTI (56–60). As hypervirulent strains become more prevalent and hypervirulent, antibiotic resistant strains emerge, it is critical to identify compartment-specific fitness factors (61, 62). Additionally, this finding that TerC is a fitness factor during urinary tract infection implies a conserved mechanism of fitness enhancement in the gut and bladder that is dispensable in the lung and blood (13, 14).

To identify this conserved mechanism, we first focused on known mechanisms of K_2_TeO_3_-induced stress that functionally overlap gut and bladder fitness factors. We determined that TerC is dispensable for metal and ROS resistance. We also determined that TerC is dispensable for the transport of several common sugars. Rather, we identified a novel role for TerC in stress tolerance, wherein Kp lacking TerC were killed more rapidly in the presence of several stresses. Given the pleiotropic effects of K_2_TeO_3_ on the bacterial cell, a role for the *ter* operon during general stress response is appealing. This could explain how the *ter* operon enhances fitness in biologically distinct sites, where the specific stresses Kp encounters are likely to differ. To identify a specific function for the *ter* operon, we undertook a systematic screen of K_2_TeO_3_ resistance in a Kp strain that lacks the *ter* operon. This screen did not reveal a consensus molecular function or specific cellular compartment of action but did suggest that K_2_TeO_3_-induced stress is primarily experienced at the cell envelope. Several of these genes were also required for growth in the presence of PmB, which is a membrane-active antibiotic. This finding affirms the necessity for TerC during stress tolerance. Finally, we demonstrate that K_2_TeO_3_ disrupts the cell envelope, though these effects were independent of *ter*, which suggests an indirect means of responding to envelope stress.

The finding that TerC enhances stress tolerance is an appealing explanation for its role in gut colonization. Host- and microbiota-derived antimicrobial peptides are critical mediators of gut homeostasis and bacterial infection (63–65). Microbiota-synthesized antimicrobial peptides are known as bacteriocins. The ability of colonizing Kp to tolerate antimicrobial peptide- and bacteriocin-mediated stress may be critical to its success in the gut when the indigenous microbiota is intact. Indeed, early experiments interrogating the biological function of the *ter* operon suggested a potential role for resisting pore-forming bacteriocins (16). Moreover, PmB is a cyclic peptide and therefore mimics the mechanism of action of many antimicrobial peptides. The necessity of TerC for tolerance to PmB suggests that TerC may also be required for tolerance to other antimicrobial peptides found in the bladder and/or gut.

Tolerance has received significant attention due to its contribution to antibiotic treatment failure and the development of antibiotic resistance; however, this neglects the role of tolerance to other stress induced by factors not characterized as classical antibiotics, such as bacteriocins or detergents. This is especially relevant in the context of PmB. Polymyxins bind to LPS in the outer and cytoplasmic membrane, resulting in cell lysis and death (55). The finding that TerC is necessary for tolerance to PmB supports the assertion proposed in other studies that the Ter proteins form a stress-sensing membrane complex that may influence transmembrane permeability (43). Similarly, the quaternary ammonium compound CPC kills bacterial cells through integration into and disruption of the cell envelope. The biological relationship between TerC and ofloxacin tolerance is less clear, though membrane peptide TisB enhances tolerance to ciprofloxacin through disruption of the proton motive force (66). Additionally, several mechanistically divergent antibiotics, including fluoroquinolones, induce inner membrane damage and cytoplasmic condensation, leading to bacterial cell death (67). Therefore, destabilization of the cell envelope may have secondary effects that impact tolerance. Exploration of TerC-dependent tolerance to other diverse stresses is likely to further refine our understanding of the biology of the *ter* operon.

Our systematic screen of K_2_TeO_3_ resistance revealed both expected and novel loci associated with K_2_TeO_3_ resistance (Table 1). Interruption of the pyruvate dehydrogenase complex resulting in increased K_2_TeO_3_ susceptibility was expected and serves as validation of the approach, as the heterologous expression of *aceE and aceF* leads to enhanced K_2_TeO_3_ resistance (68). Commensurately, TehB is a known K_2_TeO_3_ resistance protein that, in conjunction with TehA, confers resistance through the volatilization of tellurite through methyltransferase activity (18). An insertion mutant in *tehA* was not present in our Tn library. The finding that the ECA synthesis locus is a significant contributor to K_2_TeO_3_ resistance is particularly intriguing, as this conserved locus has been implicated in many critical facets of the biology of Enterobacterales. ECA is a carbohydrate structure characteristically found in the bacterial envelope, where it is attached to LPS and peptidoglycan (reviewed in (69)). Disruption in Kp reduces virulence in murine pneumonia and bacteremia models, though this phenotype is largely dependent on the stability of LPS, rather than ECA itself (70). Regardless of whether the observed K_2_TeO_3_ resistance phenotype is dependent on LPS stability, the envelope appears to be a critical mediator of K_2_TeO_3_ resistance for Kp independent of *ter*. A role for envelope stability in K_2_TeO_3_ resistance is also supported by increased K_2_TeO_3_ sensitivity of the *bamB*, *gmhB*, and *mdoG* mutants. The Bam complex (BamABCDE) is responsible for the proper insertion of proteins into the outer membrane (reviewed in (71)), GmhB plays a role in LPS biosynthesis (72, 73), and MdoGH is critical for the biosynthesis of osmoregulated periplasmic glucans (reviewed in (74)). Therefore, disruption of these genes or the ECA biosynthesis locus may result in a de-stabilized envelope.

Although this study revealed novel aspects of the biology of the *ter* operon, it is not without its limitations. First, this study provides insight into the function of the *ter* operon, but the molecular mechanisms underlying its function remain unknown. Second, the use of a transposon library that is comprised of single-gene insertions may limit the ability to identify every gene involved in K_2_TeO_3_ resistance. Some of these insertions may not sufficiently disrupt gene function to the same degree as alternative insertions present in a more complex library. Finally, experiments in this study were limited to two strains: the *ter* operon containing strain NTUH-K2044 and the *ter* operon lacking strain KPPR1. The use of these strains is convenient due to available molecular tools and because they are well-characterized; however, both strains are hypervirulent stains, and therefore do not represent the complete genomic breadth of Kp. While the *ter* operon is highly associated with hypervirulent Kp, it is not limited to hypervirulent strains (14). Interestingly, an analogous study using a murine UTI model reported similar bladder bacterial loads using a non-hypervirulent strain (75). Future studies dissecting the function of the *ter* operon should consider both hypervirulent and non-hypervirulent Kp strains. Despite these limitations, this study represents a significant advancement in our understanding of the role of the *ter* operon during Kp pathogenesis.

## MATERIALS AND METHODS

### Ethics statement

Human sample collection was approved by and performed in accordance with the Institutional Review Boards (IRB) of the University of Michigan Medical School (Study number HUM00004949). Animal studies were performed in strict accordance with the recommendations in the Guide for the Care and Use of Laboratory Animals [102]. The University of Michigan Institutional Animal Care and Use Committee approved this research (PRO00009173).

### Materials, media, and bacterial strains

All materials and chemicals were purchased from Sigma-Aldrich (St. Louis, MO) or Fisher Scientific (Hampton, NH) unless otherwise noted. The construction and validation of the isogenic *terC* mutant, the pTerZ-F complementation plasmid, and empty vector and complemented strain is described elsewhere (13, 14). All strains were grown in the presence of appropriate antibiotics for all experiments.

### Murine UTI model

The ascending UTI model used in this study has been described elsewhere (27, 76, 77). Briefly, the NTUH-K2044 and NTUH-K2044Δ*terC* strains were cultured overnight from single colonies in LB at 37° C. After overnight growth, strains were mixed 1:1 and adjusted to a final concentration of 2 x 10^9^ CFU/mL in sterile PBS. An aliquot of this inoculum was plated on LB broth containing appropriate antibiotics and plates were incubated overnight 27° C to enumerate input CFU and exact ratio. Then, mice were anesthetized with a weight-appropriate dose (0.1 ml for a mouse weighing 20 g) of ketamine-xylazine (80 to 120 mg/kg ketamine and 5 to 10 mg/kg xylazine) by intraperitoneal injection. 50 μL of inoculum was administered transurethrally into male CBA/J mice over a 30 second period to deliver 10^8^ CFU per mouse. After 48 hours, urine was collected, mice were euthanized by inhalant anesthetic overdose and the bladder was collected in sterile PBS. Bladders were homogenized, and all samples were plated on LB containing appropriate antibiotics using an Autoplate 4000 (Spiral Biotech, Norwood, MA) and incubated overnight 27° C to enumerate CFU. Bladder homogenates for growth assays were prepared from uninfected mice. Bladders were collected into 1 mL sterile PBS, homogenized, then centrifuged at 21,130 x *g* for 5 min at 4°C to pellet contaminating bacteria, then supernatant was stored at −80°C until use.

### Human urine collection

Human urine was collected from women (ages 21-40) from whom informed consent had been obtained, who have no symptoms of UTI or bacteriuria, and who had not taken antibiotics in the prior two weeks. De-identified samples from least four volunteers were pooled and filter sterilized using a 0.22-μm filter (MilliporeSigma, Burlington, MA), as previously described (77).

### Growth assays

The NTUH-K2044 pVector, NTUH-K2044Δ*terC* pVector, and NTUH-K2044Δ*terC* pTerZ-F strains were cultured overnight from single colonies in M9 minimal medium containing 0.5% glucose (M9-Glu), then diluted to OD_600_ of 0.01 in M9 minimal medium containing 0.4% or 0.5% arabinose, fucose, galactose, glucose, lactose, rhamnose, sucrose, xylose, 100% human urine, or 100% murine bladder homogenate. 100 μL of this subculture was plated into a single well of a U bottom 96-well plate in triplicate, then that plate was sealed using optical adhesive film (Applied Biosystems, Waltham, MA). This plate was incubated at 37° C with aeration and OD_600_ readings were taken every 15 min using an Eon microplate reader with Gen5 software (Version 2.0, BioTek, Winooski, VT) for 24 hours. Area under the curve was quantified using Prism 8 (GraphPad Software, La Jolla, CA).

### Minimum inhibitory concentration determination

The NTUH-K2044, NTUH-K2044Δ*terC*, NTUH-K2044 pVector, NTUH-K2044Δ*terC* pVector, and NTUH-K2044ΔterC pTerZ-F strains were cultured overnight from single colonies in M9-Glu. Overnight cultures were diluted to 10^7^ CFU/mL into M9-Glu with a 2X concentration of strain-appropriate antibiotics. Then, metals, ROS generators, antibiotics, biocides, or K_2_TeO_3_ was diluted in M9-Glu to 2X final concentration. 100 μL of this solution was plated into a single well of a U bottom 96-well plate in triplicate, then 2-fold serial diluted into M9-Glu 10 times, discarding 50 μL of the last dilution to achieve a final volume of 50 μL in each well, leaving the last well as only media. Finally, 50 μL of the 2X culture dilution was plated across each serial dilution to achieve a final cell density of 5 x 10^6^ CFU/mL. This plate was sealed using optical adhesive film (Applied Biosystems, Waltham, MA) and incubated at 37° C for 24 hours. After 24 hours, the MIC of each compound was defined as the lowest concentration that fully inhibited bacterial growth. Due to the opacity of high concentration metal solutions, MICs of metals were confirmed by replicate plating onto LB-agar and overnight culture at 37° C.

### Killing assays

The NTUH-K2044, NTUH-K2044Δ*terC*, NTUH-K2044 pVector, NTUH-K2044Δ*terC* pVector, and NTUH-K2044Δ*terC* pTerZ-F strains were cultured overnight from single colonies in M9-Glu. To ensure culture uniformity, overnight cultures were diluted 1:1,000 into fresh M9-Glu. Following overnight growth, 500 μL of culture was removed, cells were pelleted at 10,000 x *g* for 3 min, washed once in 500 μL sterile phosphate-buffered saline (PBS), then resuspended in 500 μL sterile PBS and serial plated onto LB-agar containing appropriate antibiotics to determine initial cell density. Then, K_2_TeO_3_, ofloxacin, polymyxin B, and cetylpyridinium chloride were added to these cultures to a final concentration of 1 mM, 250 μg/mL, 500 μg/mL, and 25 μM, respectively, from stocks prepared in M9-Glu. 500 μL of culture was removed at indicated time points, and cells were processed as above. All plates were incubated overnight at 27° C, and bacterial CFU/mL was quantified after overnight growth. Bacterial killing was summarized as the fraction of bacterial cells recovered at each timepoint, wherein the CFU/mL at a given timepoint was normalized to the initial CFU/mL. Area under the curve was calculated and minimum duration killing was interpolated from kill curves following log transformation of fraction recovered values using Prism 8 (GraphPad Software, La Jolla, CA).

### Tn library screen

Construction, ordering, and condensation of the KPPR1 Tn library has been described elsewhere (41, 52). This arrayed library was cultured overnight in flat bottom 96-well plates at 37° C in LB containing 40 μg/mL kanamycin. After overnight growth, arrayed Tn insertion mutants were sub-cultured into U bottom 96-well plates LB containing 40 μg/mL kanamycin and 1 μM K_2_TeO_3_ and cultured overnight at 37° C. Bacteria growth after 24 hours was measured at OD_600_ using an Eon microplate reader with Gen5 software (Version 2.0, BioTek, Winooski, VT). This assay was independently repeated twice to achieve three replicates of K_2_TeO_3_ growth. Candidate genes involved in K_2_TeO_3_ resistance were those where the mean growth in the presence of 1 μM K_2_TeO_3_ was two standard deviations above (mean OD_600_ = 0.980) or below (mean OD_600_ = 0.495) the mean of growth of all strains (mean OD_600_ = 0.738). Candidate insertion mutants and the parent strain KPPR1 were then cultured overnight from single colonies in LB containing appropriate antibiotics. Overnight cultures were diluted to 10^7^ CFU/mL into LB with a 2X concentration of appropriate antibiotics. Then, K_2_TeO_3_ was diluted in LB to 2X final concentration. Serial dilution was performed as above (see “Minimum inhibitory concentration determination”), except the 2X K_2_TeO_3_ solution was serially diluted 6 times instead of 10. 2X culture dilution was then plated across each serial dilution and the plate was sealed using optical adhesive film (Applied Biosystems, Waltham, MA) and incubated at 37° C for 24 hours. Bacterial growth was measured at OD_600_ using an Eon microplate reader with Gen5 software (Version 2.0, BioTek, Winooski, VT) and exact IC_50_ values were interpolated using a sigmoidal four-parameter logistic curve using Prism 8 (GraphPad Software, La Jolla, CA). This was repeated 3-5 times per candidate insertion mutant. Candidate insertion mutants were considered validated if their mean exact IC_50_ value was high or lower than the parent KPPR1 strain corresponding to their original screen results.

To further characterize validated insertion mutations, gene name, annotation, and BRITE Functional Hierarchies were assigned using the Kyoto Encyclopedia of Genes and Genomes (78, 79) using the VK055 gene number as the search criteria. Cellular compartment was assigned using gene ontology terms in UniProt (80). GO enrichment analysis was performed using the PANTHER Overrepresentation Test (release 2021-02-24) *Escherichia coli* as the reference list (81). For validation experiments, insertion mutants and the parent strain KPPR1 were cultured at 37° C in LB broth containing appropriate antibiotics and arrayed into flat bottom 96-well plates in triplicate. Then, arrayed insertion mutants were diluted 1:100 into LB broth containing appropriate antibiotics and 0.5 μg/mL polymyxin B in U bottom 96-well plates. Plates were sealed optical adhesive film (Applied Biosystems, Waltham, MA) and incubated at 27° C for 24 hours. After 24 hours, bacterial growth was measured at OD_600_ using an Eon microplate reader with Gen5 software (Version 2.0, BioTek, Winooski, VT). This assay was repeated twice more to achieve three replicates.

### Ethidium bromide accumulation assay

The NTUH-K2044 pVector, NTUH-K2044Δ*terC* pVector, and NTUH-K2044Δ*terC* pTerZ-F strains were cultured overnight from single colonies in M9-Glu. Following overnight growth, approximately 2 x 10^9^ CFU were harvested by centrifugation, resuspended in 2 mL of sterile PBS with or without 1 mM K_2_TeO_3_ or 500 μg/mL polymyxin B, and incubated at 37° C with shaking at 225 rpm. 1 mL of cells were immediately removed, harvested by centrifugation, resuspended in 1 mL of PBS containing 10 µM ethidium bromide, and incubated at room temperature in the dark for 10 minutes. Following incubation, fluorescence was measured in black-walled, clear-bottomed 96-well plates using an excitation of 510 nm and emission of 600 nm using a Synergy H1 Hybrid Multi-Mode microplate reader with Gen5 software (Version 2.0, BioTek, Winooski, VT). Bacterial density was measured in tandem at OD_600_. This procedure was repeated after 60 minutes of exposure to PBS with or without K_2_TeO_3_ or polymyxin B. Fluorescence was normalized to bacterial density, then fold change in relative fluorescent units was determined by dividing normalized relative fluorescent units at 60 minutes to normalized relative fluorescent units at 0 minutes.

### Statistical analysis

For *in vitro* studies, all experimental replicates represent biological replicates performed on different days. For statistical analysis, experimental values were log transformed and two-tailed ratio paired t test or RM one-way ANOVA followed by Tukey’s multiple comparison test was used to determine significant differences between groups. For *in vivo* studies, all experiments were repeated twice with independent bacterial cultures. Following CFU quantification, competitive indices ((CFU mutant output/CFU WT output)/(CFU mutant input/CFU WT input)) were calculated, then log transformed and a one-sample t test compared to a hypothetical value of 0 was used to determine significance. The limit of detection was used for the CFU output value in the case that no mutant or WT CFU were recovered in the experimental output. A P value of less than 0.05 was considered statistically significant for all experiments, and analysis was performed using Prism 8 (GraphPad Software, La Jolla, CA).

## Acknowledgements

The authors would like to thank the members of the Mobley and Bachman labs for their thoughtful feedback on this study. The authors would also like to thank the urine donors for their contribution to this study.

## Author contributions

Conceptualization: SM, JV, MAB

Methodology: JV, LAM, HLTM, MAB

Investigation: SM, JV, SNS

Visualization: JV

Funding acquisition: JV, HLTM, MAB

Project administration: MAB

Supervision: HLTM, MAB

Writing – original draft: SM, JV, MAB

Writing – review & editing: SM, JV, SNS, LAM, HLTM, MAB

## Data and materials availability

All source data for this study are provided with this manuscript.

