## Supplementary Figures for "The *Klebsiella pneumoniae ter* operon enhances stress tolerance"

**Supplementary Information**

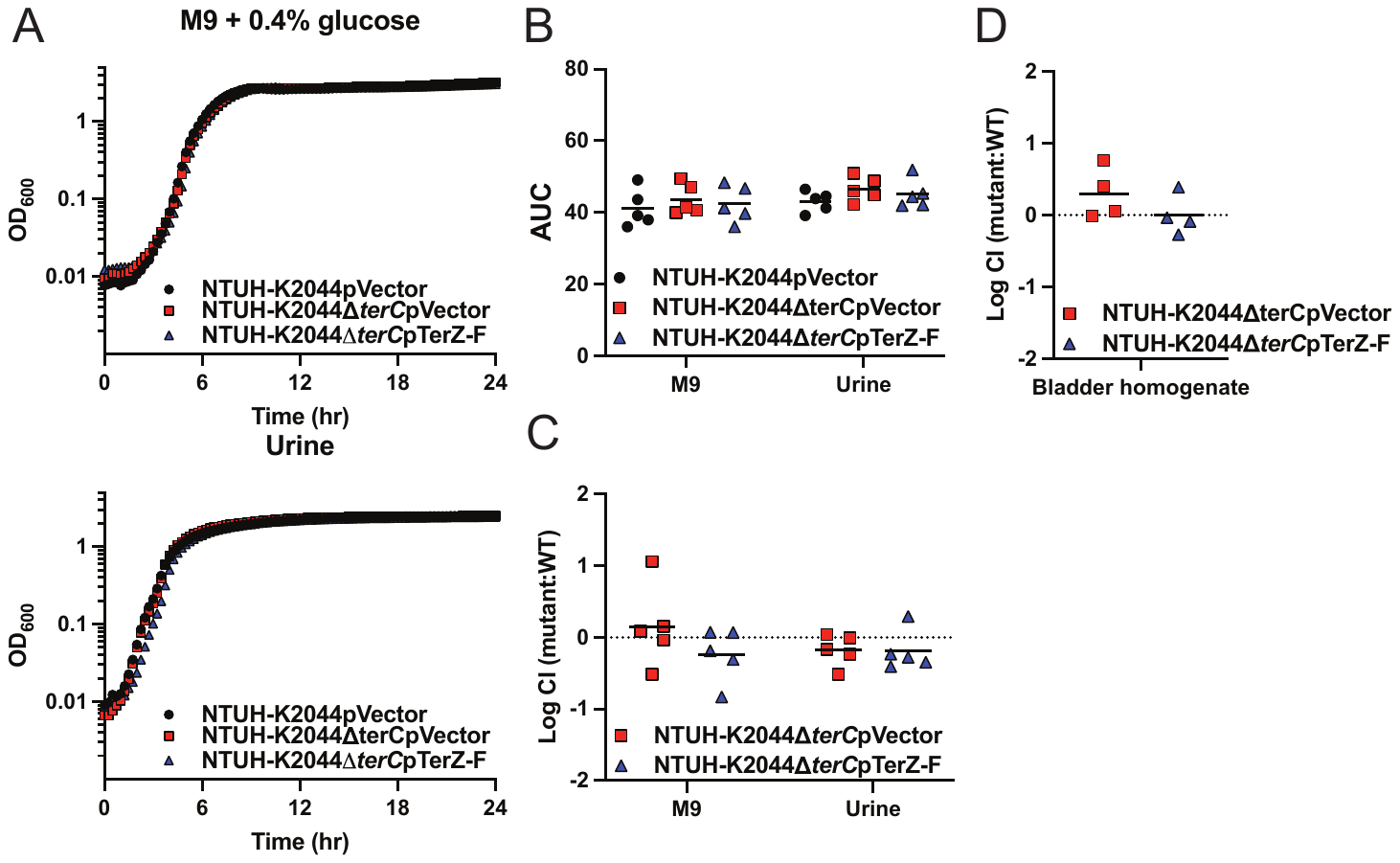
**Figure S1. TerC is dispensable for urine growth, and *ex vivo* fitness in urine and bladder homogenate**

(A) The NTUH-K2044 pVector, NTUH-K2044Δ*terC* pVector, and NTUH-K2044Δ*terC* pTerZ-F strains were grown in M9 minimal medium supplemented with 0.4% glucose or 100% human urine. (B) AUC analysis of NTUH-K2044 pVector, NTUH-K2044Δ*terC* pVector, and NTUH-K2044Δ*terC* pTerZ-F growth in M9 minimal medium supplemented with 0.4% glucose or 100% human urine (n = 5 independent experiments, Tukey’s multiple comparison test following ANOVA, mean displayed). NTUH-K2044 pVector was mixed 1:1 with NTUH-K2044Δ*terC* pVector or NTUH-K2044Δ*terC* pTerZ-F and competed in M9 minimal medium supplemented with 0.4% glucose or 100% human urine (C) or murine bladder homogenate (D). Bacterial burden was measured after 24 hours, and the competitive index of NTUH-K2044Δ*terC* pVector or NTUH-K2044Δ*terC* pTerZ-F compared to the NTUH-K2044 pVector strain was calculated for each sample (n = 5 independent experiments, mean displayed, one-sample *t*-test). For (B-C), each point represents a biological replicate, and for (D) each point represents an individual mouse bladder.

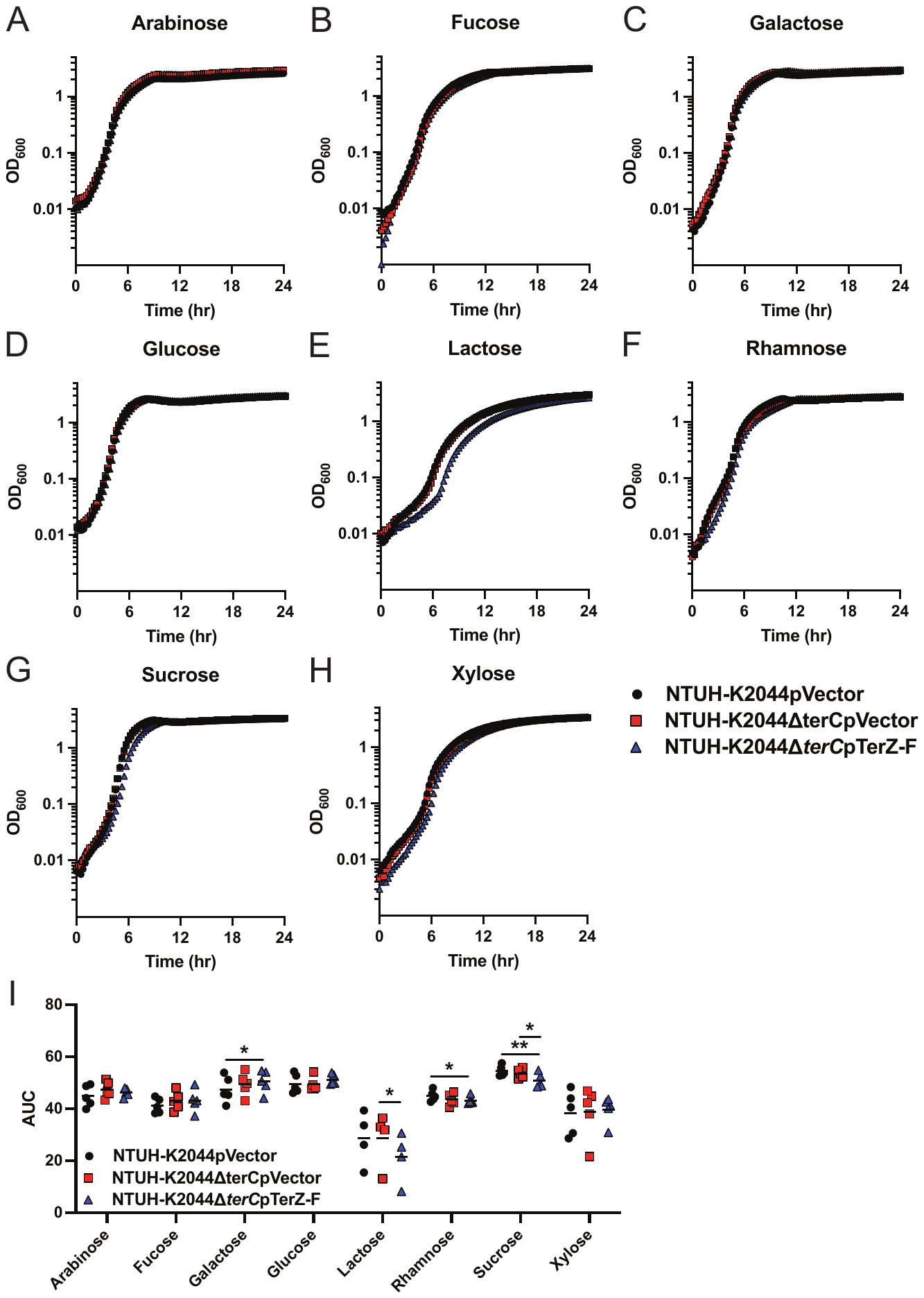

**Figure S2. TerC is dispensable for growth in several single-carbon source conditions**

The NTUH-K2044 pVector, NTUH-K2044Δ*terC* pVector, and NTUH-K2044Δ*terC* pTerZ-F strains were grown in M9 minimal medium supplemented with 0.5% (A) arabinose, (B) fucose, (C) galactose, (D) glucose, (E) lactose, (F) rhamnose, (G) sucrose, or (H) xylose. Curves are representative of four or five independent experiments. (I) Area under the curve (AUC) was calculated for each growth curve (n = 4-5 independent experiments, mean displayed, *P < 0.05, **P < 0.005, one-way ANOVA followed by Tukey’s multiple comparison test).

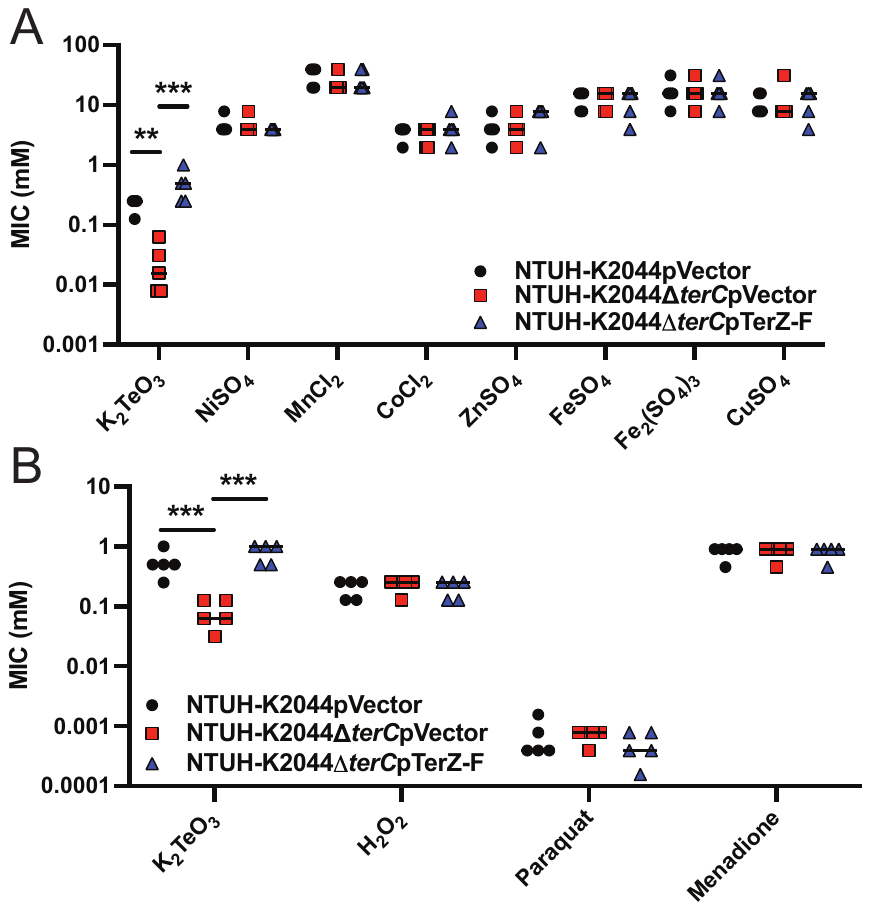

**Figure S3. TerC is dispensable for resistance to several first-row transition metals and ROS**

(A) The minimum inhibitory concentration (MIC) of several first-row transition metals for NTUH-K2044 pVector, NTUH-K2044Δ*terC* pVector, and NTUH-K2044Δ*terC* pTerZ-F was determined by broth microdilution (n = 5 independent experiments, median displayed, **P < 0.005, ***P < 0.0005, one-way ANOVA followed by Tukey’s multiple comparison test). (B) The MIC of several ROS generators for NTUH-K2044 pVector, NTUH-K2044Δ*terC* pVector, and NTUH-K2044Δ*terC* pTerZ-F was determined by broth microdilution (n = 5 independent experiments, median displayed, ***P < 0.005, one-way ANOVA followed by Tukey’s multiple comparison test).
